# Finding new cancer epigenetic and genetic biomarkers from cell-free DNA by combining SALP-seq and machine learning: esophageal cancer as an example

**DOI:** 10.1101/2020.01.18.911172

**Authors:** Shicai Liu, Jian Wu, Qiang Xia, Hongde Liu, Weiwei Li, Xinyi Xia, Jinke Wang

## Abstract

**Background:** Cancer is an important public health problem worldwide and its early diagnosis and effective prognosis are critical for its treatment. In recent years, as a good material for cancer liquid biopsy, plasma cell-free DNA (cfDNA) has been widely analyzed by next generation sequencing (NGS) for finding new molecular markers for cancer diagnosis such as size, methylation and end coordinate. However, the current studies did not still involve esophageal cancer (ESCA), a main cancer that seriously threatens human health and life in China. Here we therefore tried to find new biomarkers for this cancer from cfDNA.

**Materials & methods:** Thirty cfDNA samples from 26 ESCA patients and 4 healthy people were used to construct the NGS libraries and sequenced by using SALP-seq. The sequencing data were analyzed with variant bioinformatics methods for finding ESCA molecular biomarkers.

**Results & conclusion:** We identified 103 epigenetic markers (including 54 genome-wide and 49 promoter markers) and 37 genetic markers for ESCA. These markers provide new molecular biomarkers for ESCA diagnosis, prognosis and therapy. Importantly, this study provides a new pipeline for finding new molecular markers for cancers from cfDNA by combining SALP-seq and machine learning. Finally, by finding new molecular markers for ESCA from cfDNA, this study sheds important new insights on the clinical worth of cfDNA.

## Introduction

Cancer is an important public health problem worldwide. Its morbidity and mortality are increasing year by year, and the treatment effect is poor, which seriously affects people’s health and quality of life. According to GLOBOCAN data^1^, there were approximately 18.08 million new cancer cases and 9.56 million deaths in the world in 2018, most of whom live in low- or middle-income countries. It is estimated that by 2025, there will be about 20 million new cases of cancer every year^2^. The latest cancer statistics show that^3^, 1,762,450 new cancer cases and 606,880 cancer deaths are projected to occur in the United States in 2019. Although the incidence of cancer has not changed significantly before, cancer mortality has continued to decline, not only because of the development of medical standards, but also for preventive screening.

Tissue biopsy is still the gold standard for diagnosing tumors, but due to the traumatic nature of patients, it brings a lot of interference to the dynamic treatment of patients. There are also many risks and ethical issues in tissue biopsy, which makes it have certain limitations. Liquid biopsy is a kind of cutting-edge technology to analyze a range of tumor material in the blood or other body fluids in a minimally invasive or noninvasive manner. The tumor material includes circulating tumor cells (CTCs), cell-free tumor DNA (ctDNA), messenger RNA (mRNA), microRNA (miRNA), and exosomes. Of these tumor biomarkers, ctDNA are most widely recognized in clinical application^4^. Comparing to ctDNA, cell-free DNA (cfDNA) is a broader term which describes DNA that is freely circulating, but is not necessarily of tumor origin. CtDNA refers to the cfDNA from tumor.

CfDNA in human body fluids, such as plasma, was discovered in 1948^5^. Most of the plasma cfDNA originated from the hematopoietic system in healthy subjects, but in clinical patients (e.g. pregnancy and cancer), the related cells/tissues would release additional DNA into the plasma^6,7^. The detection of this perturbation would allow us to diagnose the abnormality for people in a noninvasive way. In recent years, methods based on the analysis of plasma cfDNA which is as an emerging technology have been largely explored for noninvasive prenatal testing (NIPT) and cancer liquid biopsy^8,9^. For example, the cfDNA based fetal aneuploidy test in pregnant women was routinely deployed in many countries by 2014, and the market value is estimated to reach 3.6 billion USD in 2019^10,11^. Studies based on cfDNA for cancer detection and tumor origin determination have also demonstrated high clinical potential^12-14^. In these studies, a variety of methods were developed for differentiating the cfDNA molecules released by the tissues-of-interest (e.g., ctDNA in cancer patients) from the background ones^15,16^. Some methods have utilized genetic biomarkers, such as the fetal-specific informative single nucleotide polymorphism (SNP) sites in pregnancies and somatic mutations in cancer patients^17,18^. However, such genetic biomarkers usually vary from case to case, which makes the development of sensitive and generalizable approaches challenging. Under this circumstance, the epigenetic biomarkers are more favored.

ESCA continues to be a leading cause of cancer death worldwide, and approximately 480,000 cases are diagnosed annually worldwide^19^. Although technological advances in surgical treatment and systemic therapy during the past few decades, over 400,000 cases have died from ESCA within the last 5 years^20^. The predicted 5-year survival rate of ESCA, which ranges from 15% to 20%, has barely improved in recent decades due to high recurrence rates, early metastatic tendency and limited knowledge of biomarkers and potential therapeutic targets^21,22^. Thus, finding new biomarkers to diagnose for ESCA is needed urgently, especially those biomarkers usable to liquid biopsy for early discovery, diagnosis and prognosis of ESCA.

In this study, we tried to find both epigenetic and genetic biomarkers of ESCA in cfDNA with adapted SALP-seq in combination with machine learning. The adapted SALP-seq^23,24^, developed by our laboratory, is a new single-stranded DNA library preparation technique. This technique is particularly suited to construct the next generation sequencing (NGS) libraries for highly degraded DNA samples such as cfDNA^24^. Moreover, by using the barcoded T adaptors, this technique is competent to analyze many cfDNA samples in a high-throughput format^24^. In this study, the NGS libraries of 20 cfDNA samples, which were from 11 pre-operation ESCA patients, 5 post-operation ESCA patients and 4 healthy people, were constructed by using SALP-seq. Based on bioinformatics analysis of the sequencing data, we identified 103 epigenetic markers (including 54 genome-wide and 49 promoter markers) and 37 genetic markers for ESCA, which may ultimately contribute to the development of effective diagnostic and therapeutic approaches for ESCA. Furthermore, these markers were verified by analyzing 10 new cfDNA samples from pre-operation ESCA patients.

## Materials and methods

### Sample Processing and Sequencing

Plasma DNA libraries were constructed from whole blood with adapted SALP^23,24^. Twenty Illumina-compatible libraries were generated. The libraries’ concentration were measured with Qubit 2.0 and mixed with equal DNA quality (ng) to generate a final sequencing library. The library was sequenced by two lanes of Illumina Hiseq X Ten platform (Nanjing Geneseeq). Paired-end sequencing was performed. The details of sample processing and sequencing protocols were described in the Supplemental Methods. In the validation experiment we sequenced 10 more cfDNA samples using the same method.

### Analysis and statistics of cfDNA sequencing data

The raw sequencing data of cfDNA were separated with the barcode by using homemade perl scripts. Then the constant sequence (19 bp) and barcode (6 bp) sequences were removed from the 5’ end of the pair-end sequencing reads 2. All sequencing reads were analyzed using the Bowtie2 tool^25^, with the parameter -X 2000 to keep the long fragments. The paired-end sequencing reads were mapped to the hg19 human reference genome in a paired-end mode, allowing one mismatches for the alignment for each end. The sequence alignment map (SAM) file was converted to BAM format using samtools^26^. Only paired-end sequencing reads with both ends aligned to the same chromosome with the correct orientation were used for downstream analysis. SNV was analyzed by samtools^26^. The annotation of SNV was performed by ANNOVAR^27^ with default setting. Functional enrichment analysis through the Database DAVID (https://david.ncifcrf.gov/)^28^ identified genes’ biological significance. P-value adjusted by Benjamini-Hochberg to <0.05 established the cut-off criteria. Reads number was calculated with bedtools^29^. The openness of 1-kb regions upstream transcription start site (TSS) in different samples was detected with DEseq2^30^, regions with p < 0.05 were selected.

In the screening of ESCA-associated important regions of the whole genome, we used the mean decrease in accuracy (MDA) method, which was calculated using the randomForest package^31^ in R (http://cran.r-project.org//). MDA represents the average decrease of classification accuracy on the OOB samples when the values of a particular feature are randomly permuted. Therefore, the permutation based MDA can be utilized to evaluate the contribution of each feature to the classification. After screening for ESCA-associated important regions, we used machine learning to classify cancer and normal samples. Due to the small sample size, classifying was performed by using support vector machine (SVM)^32^ which shows many unique advantages in solving small sample, nonlinear and high-dimensional pattern recognition problems. The gene expression data were downloaded from The Cancer Genome Atlas (TCGA) data portal (https://portal.gdc.cancer.gov/), containing RNA-seq data of 23 cancers and their corresponding normal samples. The comparison of RNA-seq datasets of selected genes between cancer and normal samples were analyzed with R scripts. The H3K27ac ChIP-seq data of TE7 ESCA cell lines was downloaded from GEO database with the accession number GSE76861^33^. The ATAC-seq data of ESCA tissues derived from donors with diverse demographic features^34^ was downloaded and the hg38 coordinates were converted to hg19 using LiftOver. All tracks were shown with UCSC genome browser.

## Results

### Clinical Specimens and SALP-seq

All procedures used in this research were performed according to the Declaration of Helsinki. This study was approved by the Ethics Committee of Jinling Hospital (Nanjing, China). All participants were recruited from the Jinling Hospital, Nanjing University School of Medicine (Nanjing, China), with informed consent. The detail information of 20 whole blood samples collected from the Jinling Hospital was listed in Table S1. We sequenced the cfDNAs from as many as 20 blood samples in two lane of Illumina Hiseq X Ten platform. Previously published sequencing data^24^ sequenced the cfDNAs from the 20 blood samples in one lane of Illumina Hiseq X Ten platform were also analyzed to investigate the esophageal cancer. We merged the sequencing data of the three lane with samtools^26^ for subsequent analysis. The data analysis in this study was, unless otherwise stated, based on the data from three lane merged. In the validation experiment we sequenced 10 more cfDNA samples using the same method to construct libraries and the detail information of samples was listed in Table S1.

### Finding cancer diagnostic/prognostic markers from chromatin accessibility of promoters

We showed the distribution of cfDNA in the whole genome by calculating and normalizing the reads density in each 1-Mb window, revealing that the distribution of different samples’ cfDNA were greatly different through the whole genome (Fig. 1A). In the previous study, we have confirmed that the NGS sequencing data can be useful for characterizing the chromatin state of different types of cfDNA^24^. In mechanism, only nucleosome-protected genomic regions can be sequenced in the NGS of cfDNA^24^. In order to identify normal samples or cancer samples viewing the signal strength of the reads distribution around TSS, we calculated the reads density of ± 5-kb region around TSSs of all human genes and calculated the average reads density using deeptools (parameter: RPKM)^35^ for 20 cfDNA samples. The results show that a peak is formed around the TSS in normal samples, and a valley is formed in the pre-operation cancer samples (Fig. 1B; Fig.S1). Moreover, we performed a principal component analysis (PCA) of reads density of ± 5-kb region around the TSSs of all human genes. As a result, the cfDNAs of pre-operation ESCA patients could be clearly distinguished from those of normal people by analyzing the cfDNA data with PCA (Fig. 1C). We also sequenced ten cfDNA samples to verify this result and got consistent results (Fig. S2). The detail information of the ten cfDNA samples collected from the Jinling Hospital was listed in Table S1.

**Fig. 1.**
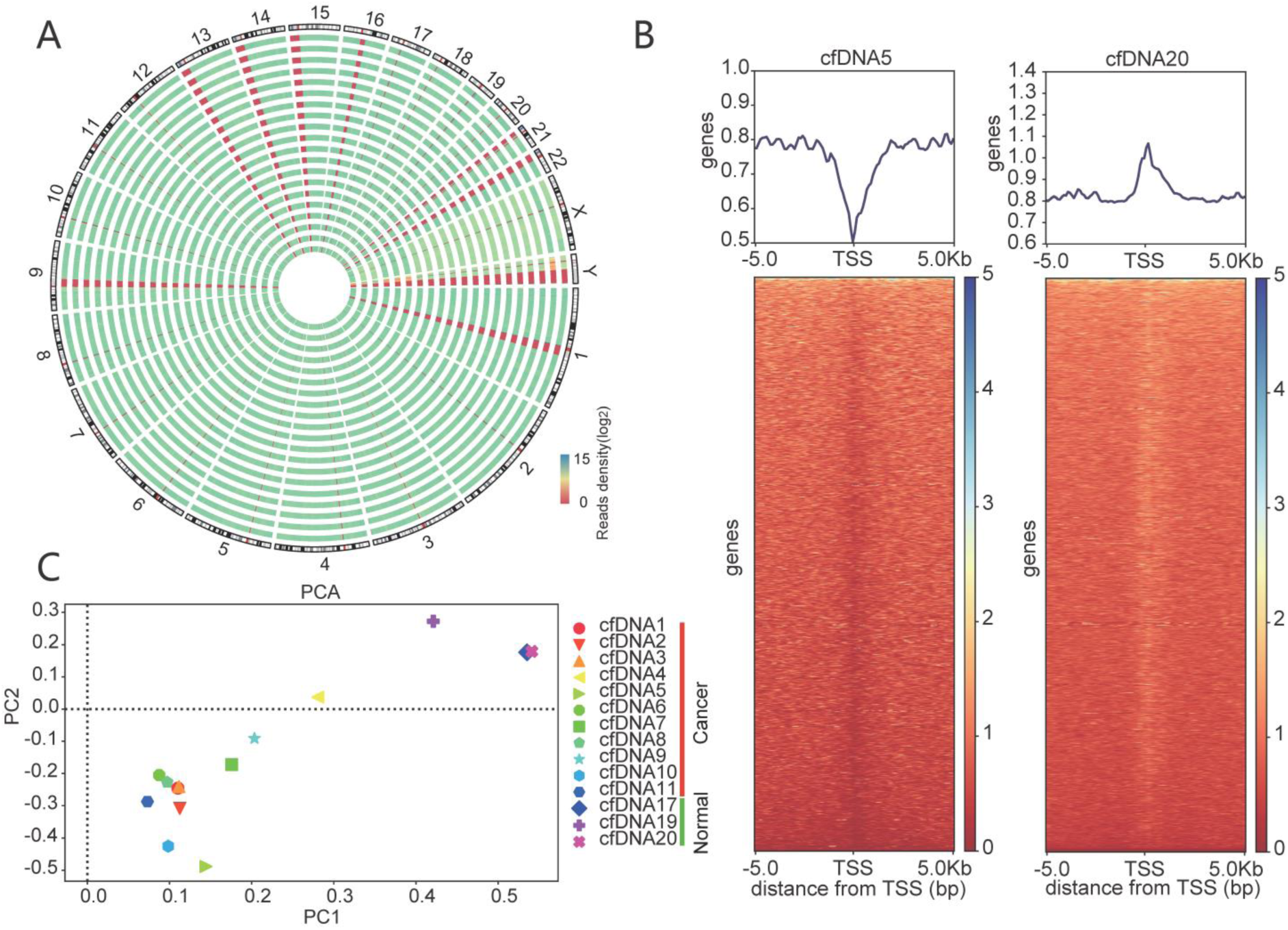
Characterization of cfDNA NGS reads distribution. (A) Distribution of reads density in each 1-Mb window of whole genome in different cfDNA samples. Reads density of cfDNA 1 to 20 were shown in order form outside to inside. (B) The signal strength of reads distribution around TSSs. The line plot shows the average signal strength of all regions around TSS. The left shows the signal strength results of cfDNA5 (cancer sample). The right shows the signal strength results of cfDNA20 (normal sample). (C) Principal component analysis (PCA) of reads density of ± 5-kb region around the TSSs of all human genes.

In the previous study, we have confirmed that the SALP-seq data of cfDNA can be useful for characterizing the chromatin state of different types of cfDNA. In this study, we differentiated cancer and normal samples by chromatin openness. The reads density of all promoters (defined as the 1–kb region upstream TSS) was calculated for each sample. The results showed that there was a great difference of reads density of promoters between normal cfDNA samples and ESCA cfDNA samples (Fig.2A). Notably, some promoters showed extremely low reads density in all cancer samples but high density in normal samples (Fig.2B). There were 49 genes had such a distinct feature. Through the clustering results of these 49 genes from the heatmap, it was clearly seen that the cancer samples can be distinguished from the normal samples (Fig. 2B). Moreover, we sequenced ten new cfDNA samples to verify this result and got consistent results (Fig. 2D). We calculated the reads density of promoters of the 49 genes in the post-operation cancer samples. The results showed that the reads density of promoters of most of these genes was significantly increased in the post-operation cancer compared with pre-operation cancer (Fig. 2C), indicating the effect of surgery. These data suggested that the chromatin accessibility of the promoters of these 49 genes can be used as the diagnostic and prognostic markers for cancer.

**Fig. 2.**
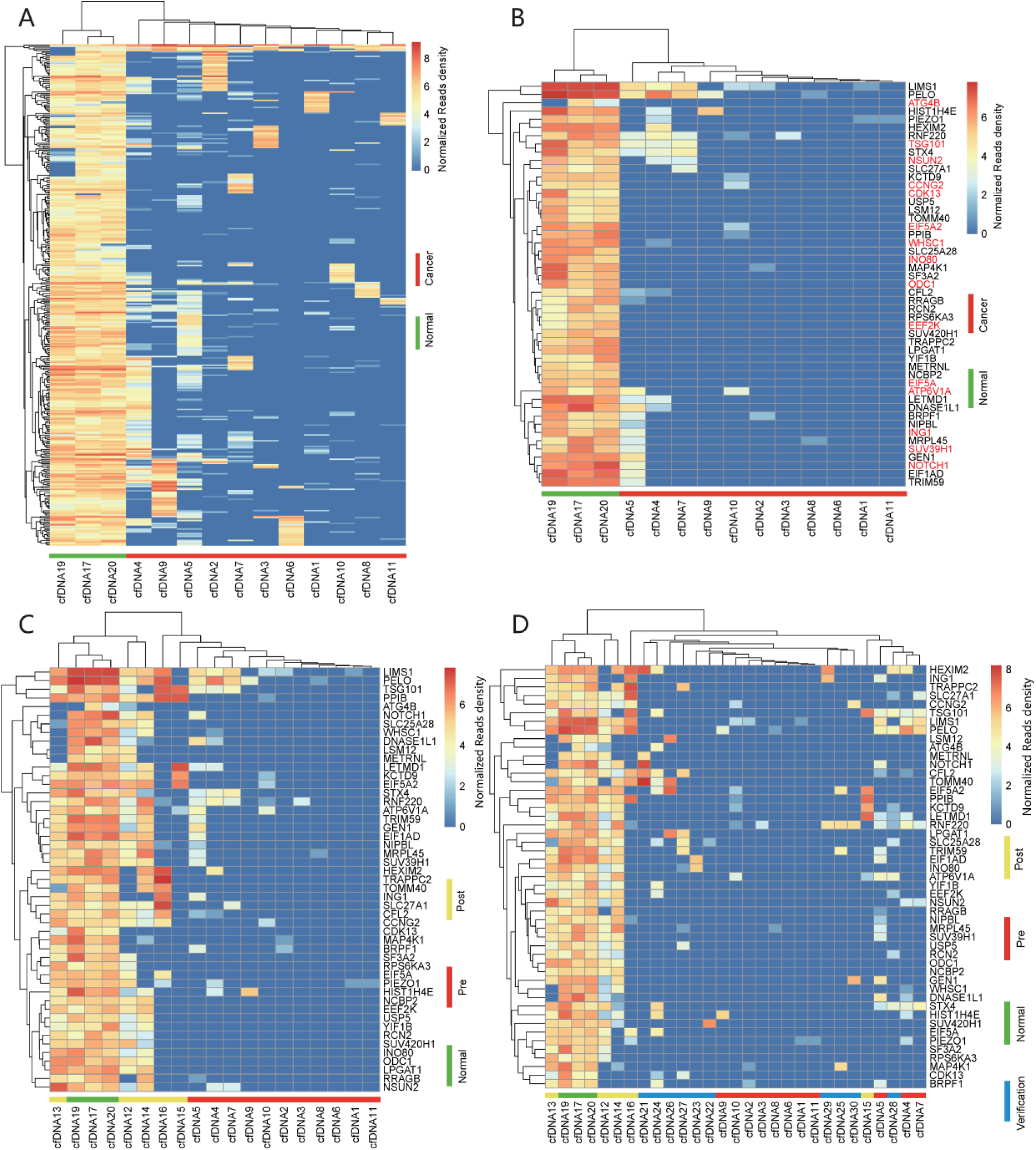
Analysis of gene promoters. (A) The heat map and clustering of reads density of promoters with significant difference in chromatin accessibility in cfDNA samples. (B) The heat map and clustering of reads density of promoters of 49 selected genes. The genes in red are known ESCA-associated genes. (C) The heat map and clustering of reads density of promoters of 49 selected genes in the post-operation cancer cfDNA samples. (D) The heat map and clustering of reads density of promoters of 49 selected genes in the 10 verification cancer cfDNA samples. Pre: pre-operation cfDNA; Post: post-operation cfDNA; Normal: normal cfDNA; Verification: 10 cfDNA samples of verification.

Through database and literature search, we found that 15 (genes in red in Fig. 2B) of these 49 genes have been reported to be closely associated with ESCA (Table S3). Therefore, we inferred that the remaining 34 genes are newly discovered genes associated with ESCA. To validate the relationship between these 49 genes and ESCA, we downloaded the RNA-seq data (containing 163 ESCA and 11 normal samples) of ESCA from the TCGA database for analysis. We found that the expression of most of these genes were significantly up-regulated in cancer samples (Fig. S3A). In the process of database and literature search, we found that some of these 49 genes are not only related to ESCA, but also associated with other cancers (Table S3). We downloaded RNA-seq data of 23 cancers from the TCGA database to analyze the relationship between these genes and various cancers. The results showed that most of these genes were significantly up-regulated in various cancers (Fig. S3B). The above results indicate that the chromatin accessibility of promoters of the selected 49 ESCA-associated genes can distinguish between normal and cancer, and chromatin accessibility of these promoters can be also used to diagnose and prognose cancer. To further understand the functional roles of these genes in ESCA, GO analysis were performed. As a result, both chromosome organization (GO: 0051276) and chromatin organization (GO: 0006325) were significantly enriched, implying that some of these genes play critical roles in regulating the chromosome or chromatin structure (Fig. S4A; Table S4). In the GO term of chromosome organization, there were nine genes, including BRPF1, NIPBL, GEN1, SUV39H1, HIST1H4E, INO80, PELO, WHSC1, and ING1. In the GO term of chromatin organization, there were seven genes including BRPF1, NIPBL, SUV39H1, HIST1H4E, INO80, WHSC1, and ING1. Among the genes enriched in GO: 0051276 and GO: 0006325, SUV39H1, INO80, WHSC1, and ING1 were known ESCA-associated genes (Table S3). Other enriched GO terms were related to regulation of cell cycle (GO: 0051726), growth (GO: 0040007) and so on, which all play an important role in cancer development (Fig. S4A; Table S4). Pathway annotation was used to screen out the altered biological functions arising from the 49 selected genes. The results indicated that these genes were mainly enriched in 5 pathways, including Lysine degradation, mTOR signaling pathway, mRNA Surveillance pathway, Insulin resistance, and Spliceosome (Fig. S4B; Table S5).

### Finding cancer diagnostic/prognostic markers from chromatin accessibility of whole genome

To find ESCA-associated important regions from the whole genome, we calculated the reads density in each 1-kb window of the whole genome, and then used MDA to screen out 88 ESCA-associated regions, most of which were non-coding sequences, suggesting that these regions contain important regulatory elements (Fig. 3). Then we pinpointed the genomic location of these regions, and found that 36.36% were located in the distal intergenic regions (more than 10-kb from TSSs), and 25% were located in the Proximal regulatory region (10-kb regions upstream TSSs) (Fig. 3, inset). It can also be seen from the MDA diagram that distal elements (defined as occurring inside of distal intergenic) is much more important than promoter elements (defined as occurring inside of Proximal regulatory region) in the classification (Fig. 3), indicating that distal elements exhibited a greater specificity and wider dynamic range of activity in association with cancer, whereas promoter element accessibility was less cancer-specific. This functional specificity of distal regulatory elements was also previously observed in healthy tissues and in cancer^34,36,37^.

**Fig. 3.**
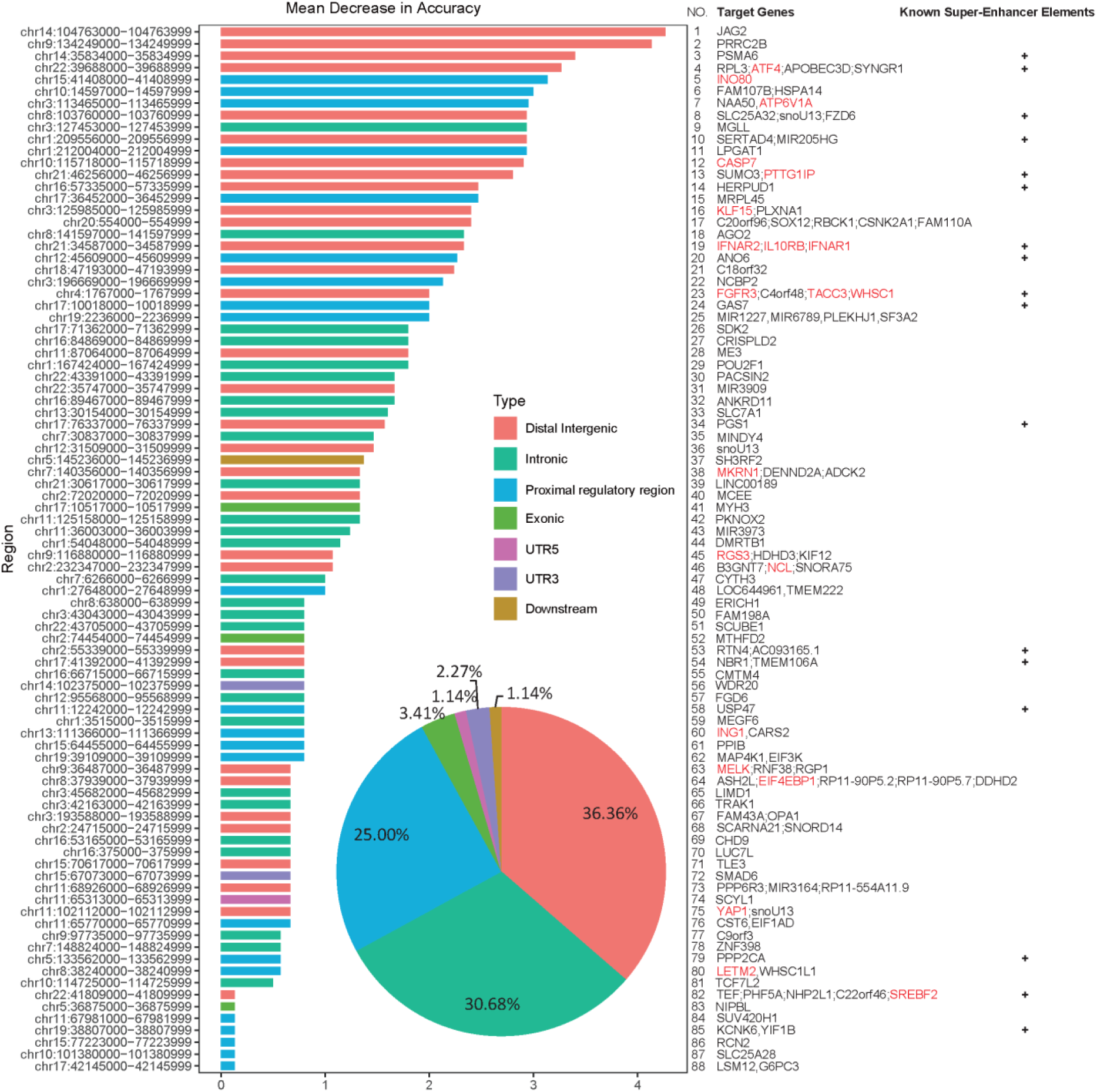
Analysis of ESCA-associated important regions in genome. The 88 ESCA-associated important regions selected based on MDA. The ordinate represents the regions, and the abscissa represents the importance value. Different colors indicate the genomic location of these regions. Inset shows the distribution of genomic location of these ESCA-associated important regions in genome. Most of the regions were located in distal intergenic, intronic, and proximal regulatory region. The genes and known super-enhancer elements assigned to these regions are shown. The genes in red are the know ESCA-associated genes.

We researched the ESCA-associated important regions that were located in distal intergenic and proximal regulatory region. The Fig. 4A showed the reads density in ESCA-associated important regions (including 32 distal intergenic and 22 proximal regulatory region) of each sample, suggesting that there was a great difference between normal cfDNA samples and ESCA cfDNA samples. In addition, we also calculated the reads density of the 54 regulatory regions in the post-operation cancer samples. The results showed that the reads density of most of these regions was increased in the post-operation cancer compared with pre-operation cancer (Fig. 4B), showing the effect of surgery. These data suggested that the chromatin accessibility of these genomic regions can be used as the diagnostic and prognostic markers for cancer. To further validate these markers, we sequenced ten new cfDNA samples to verify this result. As a result, all the consistent results were obtained (Fig. 4C). These data also indicated that more new caner-associated markers could be identified using the whole-genome chromatin accessibility characterized with cfDNA.

**Fig. 4.**
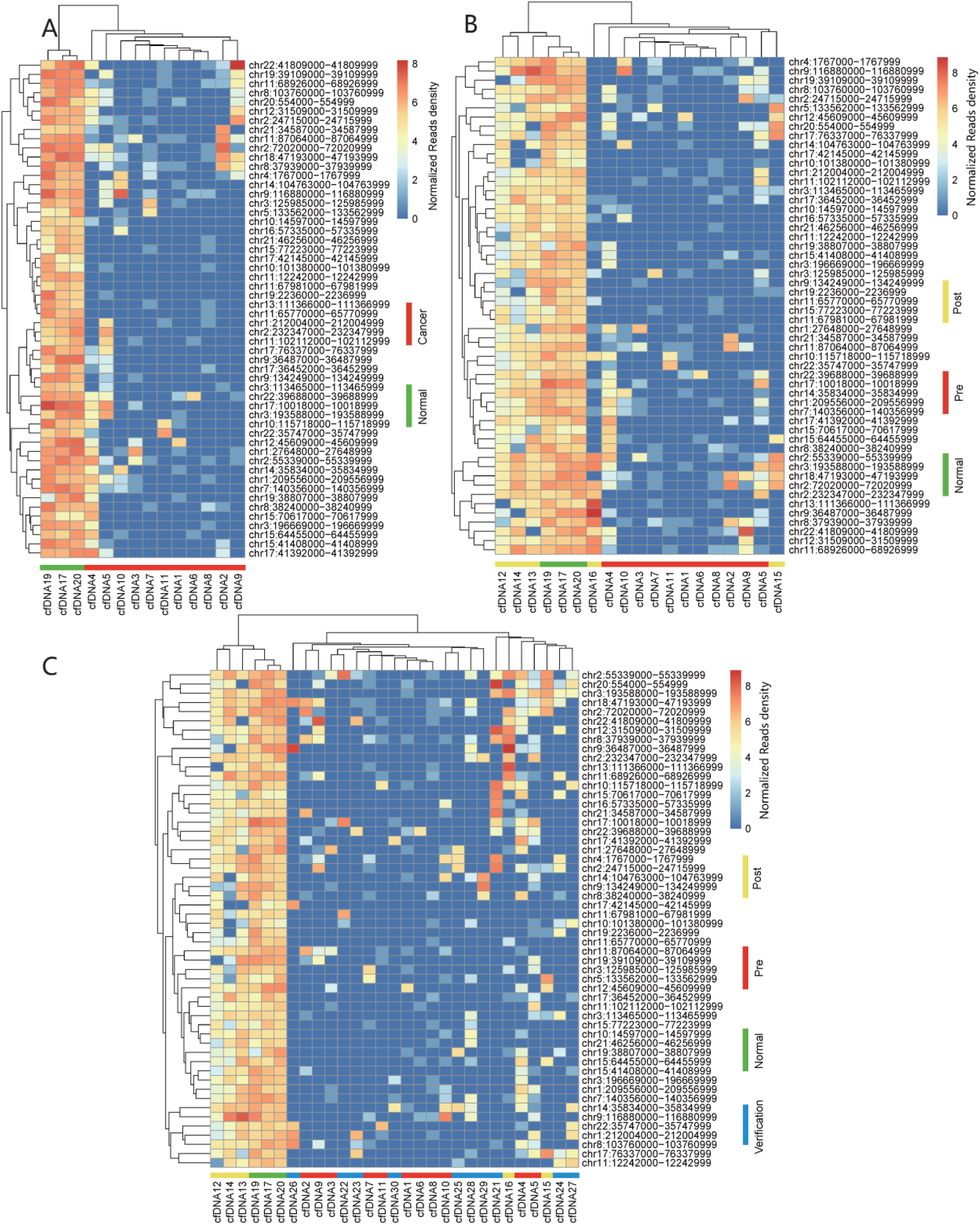
Analysis of the selected 54 ESCA-associated important regions. (A) The heat map and clustering of reads density of ESCA-associated important regions (including 32 distal intergenic and 22 proximal regulatory regions) of each sample. (B) The heat map and clustering of reads density of ESCA-associated important regions in the post-operation cancer cfDNA samples. (C) The heat map and clustering of reads density of ESCA-associated important regions in the 10 verification cancer cfDNA samples.

To further verify the chromatin accessibility of these genomic regions characterized with cfDNA using SALP-seq, we downloaded the H3K27ac ChIP-seq data of TE7 ESCA cell lines from GEO with the accession number GSE76861^33^, downloaded the ATAC-seq data of 19 ESCA tissues derived from donors with diverse demographic features^34^ from the TCGA database. These data together with SALP-seq data were compared by visualizing these regions (including 32 Distal Intergenic regions and 22 Proximal regulatory regions) with UCSC genome browser. The results revealed that the chromatin accessibility of these genomic regions characterized with cfDNA using SALP-seq in this study were highly consistent with those characterized with cancer cells and tissues using H3K27ac ChIP-seq and ATAC-seq (Fig. S5).

Because these regions are non-coding regions and their chromatin accessibility are significantly changed, they should played regulatory functions in tumor by providing accessible binding sites to transcription factors (TFs). We therefore searched the potential TF binding sites in these regions using FIMO^38^ with the motif matrix obtained from the HOCOMOCO (version 11)^39^ with default setting. The results showed that these regions contain a large number of TF binding sites (TFBSs) (Table S6). Moreover, we compared these regions with the SEdb database^40^, a comprehensive human super-enhancer database. The results revealed that 17 of these 54 regions were well known super-enhancer elements (Fig.3; Table S7).

In order to find the target genes regulated by the selected ESCA-associated distal elements and promoter elements, we predicted target genes for these genomic regions by using EnhancerAtlas^41^ with parameter “Esophagus”. The results showed that these regions were assigned to 104 genes (Fig.3; Table S7), 16 of which were also present in the 49 genes identified above with promoter chromatin accessibility, including LSM12, MAP4K1, SF3A2, SLC25A28, RCN2, YIF1B,NCBP2, EIF1AD, LPGAT1, WHSC1, SUV420H1, ATP6V1A, INO80, PPIB, MRPL45, and ING1. These data not only illustrated the reliability of our results, but also indicated that more cancer-associated genes could be identified using the whole-genome chromatin accessibility characterized with cfDNA. To further validate these potential cancer-associated genes, we downloaded the RNA-seq data of ESCA from the TCGA database for analysis. The results indicated that the expression of most of these genes were significantly up-regulated in cancer samples (Fig. S6). Through database and literature search, we found that 21 of these 104 genes have been reported to be closely associated with ESCA (Fig.3; Table S8). Therefore, we inferred that the others are newly discovered genes associated with ESCA. The gene annotation revealed that these genes were mainly associated with the biological processes of apoptotic, metabolic, cell growth, and translational initiation, and molecular function of histone lysine N-methyltransferase activity and interferon activity (Fig. S7A; Table S9). The Pathway annotation showed that these genes were mainly enriched in 9 pathways, including signaling pathways of PI3K-Akt, AMPK, Hippo, and Jak-STAT (Fig. S7B; Table S10). These biological processes, molecular functions and pathways all play important roles in cancers.

### Establishing classification model of ESCA based on the identified ESCA-associated regions

To build a classification model for predicting ESCA, we then analyzed 88 regions with the SVM algorithm^32^. The results revealed that the established SVM model could accurately distinguish cancer samples from normal samples with a AUC value of 1.0 (Fig. 5A). To further improve the clinical applicability of the classification model, after debugging and screening, we finally selected top 24 ESCA-associated important regions to re-establish the model. As a result, the re-established model could still accurately distinguish cancer and normal samples with a AUC value of 1.0 (Fig. 5B). The validation of the model with the later sequenced cfDNA samples obtained good prediction results with accuracy 93.8% (Fig. 5C). In order to explore the effect of the reads number on the model, we selected 10^6^ and 10^7^ reads from 20 firstly sequenced cfDNA samples and then predicted separately. The results showed that the model still maintained a good predictive effect at 10^7^ reads (Fig. 5D-E), suggesting that the model can be used to predict ESCA at lower cfDNA sequencing depths.

**Fig. 5.**
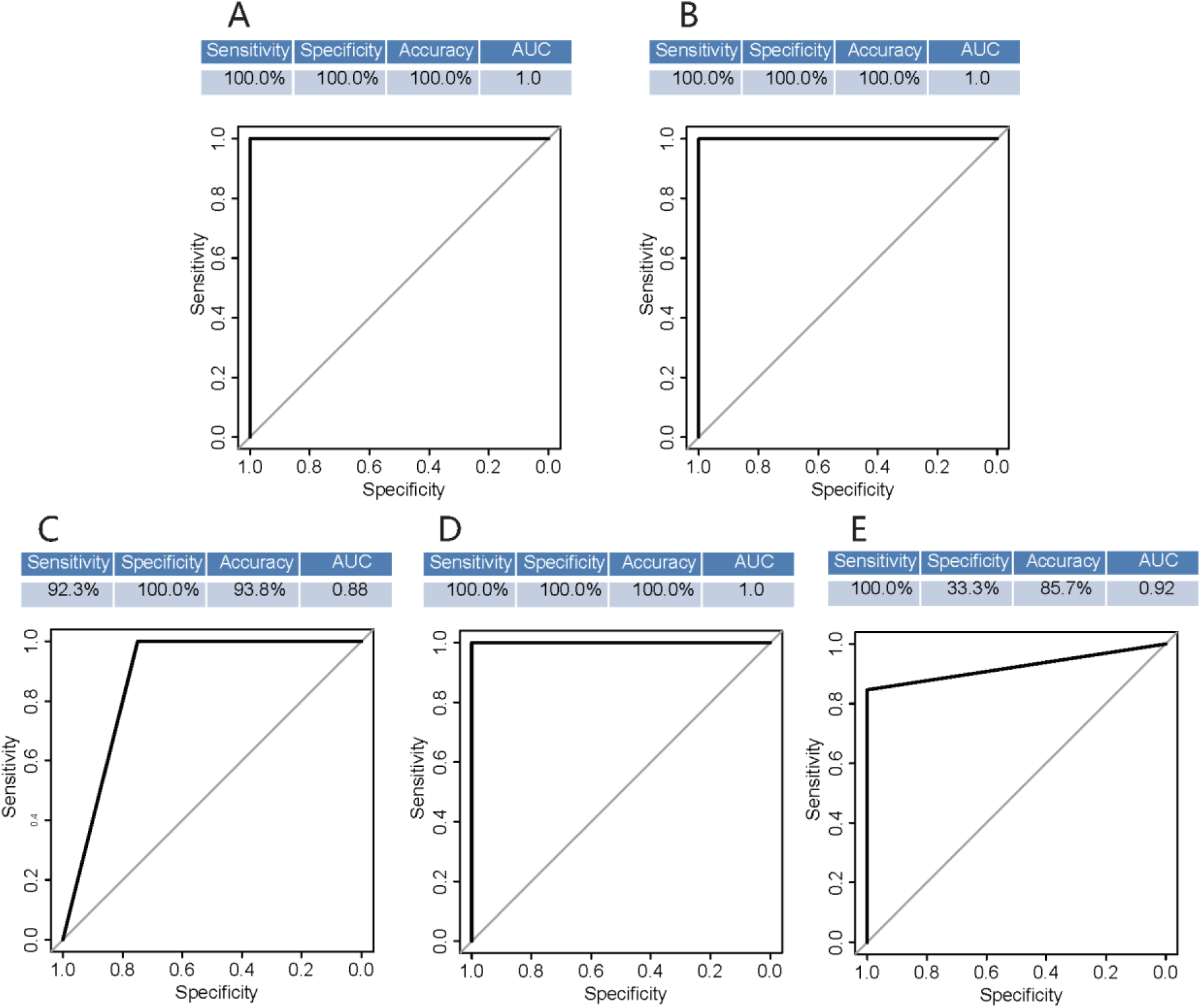
Classification of ESCA and normal cfDNA based on ESCA-associated important regions. (A) Results of classification of cancer and normal cfDNA samples using SVM based on 88 ESCA-associated important regions. (B) Classification results based on top 24 ESCA-associated important regions. The picture below shows the receiver operating characteristic curve (ROC).(C) Classification results of the verify data. (D) Classification results of 10^7^ reads which were extracted from each sample. (E) Classification results of 10^6^ reads which were extracted from each sample. The pictures of receiver operating characteristic curve (ROC) were shown.

### Characterizing ESCA-associated mutations with cfDNA

Mutations analysis of cfDNA with target or genome scale sequencing was widely used in NIPT or liquid biopsy. We next analyzed ESCA-associated mutations with SALP-seq reads of cfDNA samples. The results indicate that there are mutations in the whole genome (Fig. 6A). We extracted mutational signatures of 20 cfDNA samples using 6 kinds of base substitutions (C>A, C>G, C>T, T>A, T>C, and T>G). The results indicate there are variant levels of all these base substitutions in different individuals (Fig. 6B). The results show that the pre-operation ESCA cfDNA has lower C>T and T>C transitions than normal cfDNA, but has higher C>G and C>A transversions than normal cfDNA (Fig. 6C). There are also significant differences of Ti and Tv frequencies between pre-operation ESCA and normal samples (Fig. 6D). Moreover, there is a significant difference of Ti/Tv ratio between pre-operation ESCA and normal samples (Fig. 6E). Importantly, the surgical treatment evidently changed the seven mutation features (Fig. 6C–E). The seven mutation features can be developed into diagnostic markers for ESCA liquid biopsy. To pinpoint the genomic location of the SNV, we systematically annotated the SNV using ANNOVAR^27^. The results indicate that most of the SNVs are located in distal intergenic and intronic regions (Fig. 6F).

**Fig. 6.**
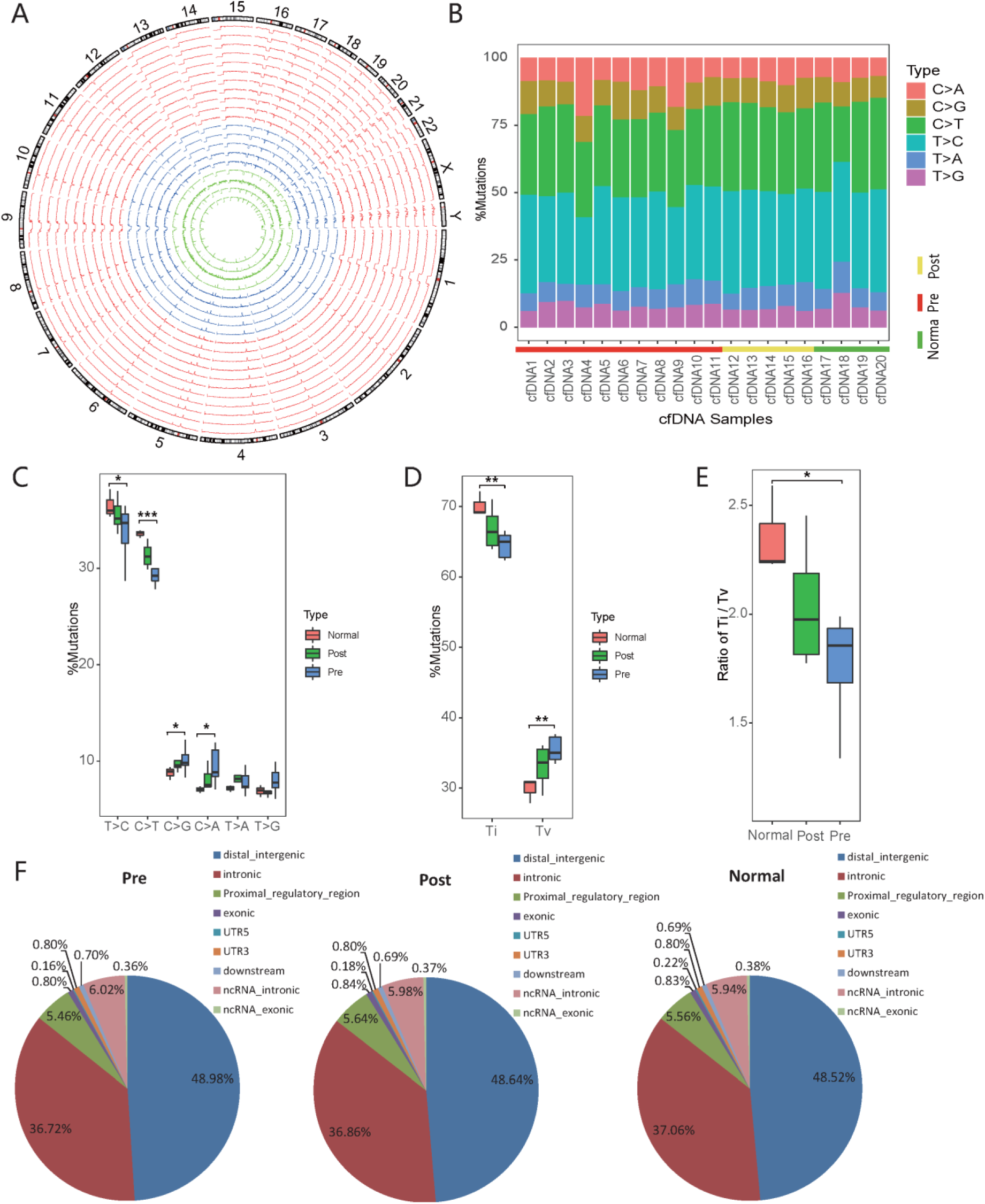
Analysis of mutations. (A) Distribution of mutation density in each 1-Mb window of whole genome in different cfDNA samples. (B) Stacked bar plot shows distribution of mutation spectra for every cfDNA sample. (C) Box plot summarizing the SNV of different types of cfDNA samples. P values (*p < 0.05, **p < 0.01, ***p < 0.001) (D) and (E) Box Plot created by dividing the SNV into Ti and Tv. Ti, transition; Tv, transversion. (F) Distribution of genomic location of the SNV in genome. Most of the SNVs were located in distal intergenic and intronic regions.

To find the mutations in coding sequences of all genes, we analyzed mutations in each cfDNA samples. The results indicate that there are large amount of mutations in thousands of genes in pre- and post-operated ESCA cfDNAs and normal cfDNA (Table S11). To test whether the clinically relevant mutations can be detected by the cfDNA NGS, we compared these genes identified by cfDNA sequencing with the MSK-IMPACT panel genes (468 genes). MSK-IMPACT can be used to identify clinically relevant somatic mutations, novel non-coding changes, and mutational features shared between common and rare tumor types, which authorized by the Food and Drug Administration of USA in 2017^42^. Finally, we found that 37 mutated genes uniquely existed in pre-operation patients (Fig.S8A; Table S12), suggesting that these genes might play a certain role in ESCA. These 37 genes contained many well-known cancer-related genes, such as PTEN, MYC, EZH1, IDH2, AKT2, and FGFR2 (Table S12).

We then performed functional enrichment analysis on these 37 genes. GO analysis reveals that these genes are significantly associated with cell death regulation, transcriptional activator activity, RNA polymerase II transcription regulatory region sequence-specific binding, tissue development and so on (Fig. S8B; Table S13).The KEGG pathway analysis demonstrates that these genes are significantly enriched in Pathways in cancer, Signaling pathways regulating pluripotency of stem cells, Central carbon metabolism in cancer, Transcriptional misregulation in cancer, and mTOR signaling pathway (Fig. S8C; Table S14). The GAD DISEASE CLASS analysis demonstrated that these genes are significantly enriched in CANCER, REPRODUCTION, and DEVELOPMENTAL (Fig. S8D; Table S15). Moreover, the GAD DISEASE analysis reveals that these genes are significantly associated with esophageal cancer (Fig. S8D; Table S15). The UP KEYWORDS analysis of DAVID demonstrates that these genes are significantly enriched in Disease mutation, Transcription regulation, Tumor suppressor and Proto-oncogene (Fig. S8E; Table S16). The UP SEQ FEATURE analysis of DAVID demonstrates that these genes are significantly enriched in mutagenesis site and sequence variant (Fig. S8E; Table S16).

## Discussion

In this study, we performed the NGS library construction and sequencing of cfDNA samples with SALP-seq that was developed by our laboratory^23^. As a new single-stranded DNA library preparation and sequencing technique, SALP-seq is particularly suited to construct the NGS libraries for highly degraded DNA samples such as cfDNA^23,24^. Moreover, by using the barcoded T adaptors, this technique is competent to analyze many cfDNA samples in a high-throughput format^24^. In this study, we used the SALP-seq to analyze as many as 30 cfDNA samples successfully. Most of samples obtained over 80% mappable reads (Table S1). Both ESCA-associated epigenetic and genetic biomarkers were successfully identified by analyzing the obtained sequencing data. The results demonstrated that this technique can be reliably applied to the future cfDNA NGS researches.

This study sheds important new insights on the clinical worth of cfDNA. In this study, we analyzed the cfDNA NGS reads data with machine learning algorithms. The results indicated that the main ESCA-associated epigenetic and genetic markers could be effectively identified by comparing the cancer and normal cfDNA samples. Especially, the cfDNA samples could be clearly classified by using the identified epigenetic markers (Fig. 2 and Fig. 4). Moreover, the promoter and genome-wide markers obtained highly consistent classification results (Fig. 2 and Fig. 4), indicating the reliability of these epigenetic markers in dicriminating cfDNA samples. Importantly, the SVM model established with the epigenetic markers could be used to accurately distinguish cancer cfDNA samples from normal cfDNA samples with a high AUC value, even by using as few as 24 most important epigenetic markers at low cfDNA sequencing depth (10^7^ reads/sample) (Fig. 5). These results reveal that important application of cfDNA in the systematic finding of cancer-associated markers, especially epigenetic markers associtated with chromatin accessibility, at the genome-wide scale.

This study provides a new pipeline for finding new molecular markers for cancers from cfDNA by combining SALP-seq and machine learning. In recent years, as a good material for cancer liquid biopsy, plasma cfDNA has been widely analyzed by next generation sequencing (NGS) for finding new molecular markers for cancer diagnosis such as fragment size^43,44^, methylation^45-47^, and end coordinate^48,49^. However, the size-based plasma DNA diagnostics still faced some limitations that may challenge its wide application^50^. The methylation detection has to do cell-free methylated DNA immunoprecipitation and high-throughput sequencing (cfMeDIP-seq)^45^. In comparison with these reported cfDNA-based methods, our method is more simple and easy. Only two steps are needed. One is SALP-seq and the other is machine learning. Our method needs no pre-treatments to cfDNA such as size selection, target enrichment, chemical treatment (e.g. bisulfite conversion), and immunoprecipitation, which not only avoid the introduction of more artificial biases in cfDNA analysis, but also greatly simplify the detection process. In the pipeline developed by this study, the cfDNA NGS reads data were analyzed with machine learning, this is different from the above mentioned previous studies. These studies did not employ machine learning to treat the NGS data. The results obtained by this study indicate machine learning can play important roles in cfDNA NGS data analysis.

By using post-operation cfDNA samples, this study showed that the identified ESCA-associated epigenetic and genetic markers are tumor-associated. In other words, these markers should come from tumor but not from other tissues such as leukocytes, because these markers changed in response to surgical operation. It was found that most of these markers became disappeared after surgery (Fig. 2D and Fig. 4C), allowing the patient’s cfDNA was classified into or near normal cfDNA (Fig. 2D and Fig. 4C). If these markers come from other tissues such as leukocytes, they should have no such evident response to surgery. Therefore, this study revealed that these markers not only come from tumor, but also are beneficial for cancer prognosis. In other words, these markers can be used to judge and track the effects of cancer treatment noninvasively. In the later visiting, five patients provided the post-operation cfDNA samples still survive at present after the surgery in 2017, suggesting the good prognosis with these markers.

This study sheds important new insights on the potential regulatory and molecular mechanisms of tumorigenesis of ESCA. By analyzing and comparing the cfDNA NGS data, this study identified 49 ESCA-associated promoters and 88 ESCA-associated genome-wide regions. These ESCA-associated chromatin regions are all non-coding DNAs. Importantly, all these regions became more accessible in ESCA, suggesting that these regions play critical regulatory roles in tumorigenesis of ESCA. Especially, many of these ESCA-associated regions are distal intergenic (32, 36.36%) and proximal regulatory regions (22, 25%) (Fig. 3). Moreover, by comparing these regions with a comprehensive human super-enhancer database, the SEdb database^40^, 17 of these 54 regions were well known super-enhancer elements (Table S7). Additionally, it was found the SVM model established with the 24 most important regions could still be used to accurately distinguish cancer samples from normal samples with a high AUC value. In these 24 regions, there are 14 distal intergenic regions (58.3%), 8 proximal regulatory regions (33.3%), and only 2 intronic regions (8.3%). By comparing with the chromatin accessibility level characterized by the H3K27ac ChIP-seq of TE7 ESCA cell lines and the ATAC-seq of ESCA tissues^33,34^, it was found that the chromatin accessibility of these regions characterized by cfDNA is highly consistent with those characterized by other methods in ESCA cells and tissues (Fig. S5). Additionally, these regions contain a large number of TFBSs (Table S6). Therefore, these non-coding ESCA-associated chromatin regions should play critical regulatory roles in tumorigenesis of ESCA. By assigning these regions to genes, 153 (49 plus 104) ESCA-associated genes were identified, in which 104 and 49 genes are connected with the genome-wide regions and promoters, respectively, and 16 are connected with both regions. It was found that 15 of 49 genes and 21 of 104 genes have been reported to be closely associated with ESCA (Fig.2B and Fig.3; Table S3 and S8). For example, studies have shown that WHSC1 has oncogenic activity and can cause protein lysine methyltransferases dysregulated in ESCA and other cancer^51^. Targeting WHSC1, currently developing specific inhibitors for diagnose and treat cancer, such as MCTP39 and LEM-06, which are in preclinical trials^51^. Furthermore, elevated expression of WHSC1 is often observed in many types of human cancers, and expression product of WHSC1 is essential for the growth of cancer cells^52-54^. The gene EIF5A2 was related to not only ESCA^55-57^, but also breast cancer^58^, lung cancer^59,60^, bladder cancer^61,62^, stomach cancer^63-65^, oral cancer^66^, liver cancer^67^, and colorectal cancer^68^. By checking the expression of these genes detected in tumors (RNA-seq data in TCGA), most of these genes are significantly up-regulated in tumors (Fig. S3 and Fig. S6), in consistent with the increased chromatin accessibility of these regions revealed by cfDNA in this study. These results indicate that these ESCA-associated regions play regulatory roles in ESCA (Fig. S3A and Fig. S6) and other cancers (Fig. S3B). The gene annotation also revealed that these genes are also closely related to ESCA and other cancers (Fig. S4 and Fig. S7). Therefore, most of genes identified by this study are newly discovered genes associated with ESCA. For example, the newly discovered ESCA-associated gene, JAG2, plays a role in NOTCH signaling and Hedgehog signaling^69-71^. Dysregulation NOTCH signaling and Hedgehog signaling are both closely related to the development of cancer (e.g. ESCA)^70,72^, further explaining the correlation between JAG2 and ESCA. The TCGA RNA-seq data revealed that JAG2 was significantly up-regulated in ESCA tissue (Fig. S6). Importantly, the JAG2-connected region identified in this study has the highest importance value of MDA (Fig. 3). Thus, targeting JAG2 may offer a promising therapeutic strategy for ESCA treatment. And the others could be potential targets for ESCA diagnosis and treatment.

Mutation of the cfDNA samples was analyzed in this study. The C>T and T>C transitions were the major SNVs (Fig. 6B). The C>T transitions may arise by replication of uracil generated by APOBEC cytidine deamination^73,74^, while the cause of T>C currently is no clue. The C>G, C>A, and potentially additional C>T substitutions may be introduced by error-prone polymerases following uracil excision and generation of abasic sites by uracil-DNA glycosylase (UNG) ^73,74^. Further analysis revealed that the differences between pre-operation ESCA samples and normal samples reach statistically significant level in the frequencies of Ti and Tv (Fig. 6D). Moreover, there was significant difference in the frequencies of Ti/Tv ratio between pre-operation ESCA samples and normal samples (Fig. 6E). These mutation characteristics of post-operation ESCA samples were closer to the normal samples, although they did not reach the same level (Fig. 6C-E). These results revealed that seven features, including the frequencies of C>T, T>C, C>G, C>A, Ti, Tv, and Ti/Tv ratio, could be developed as diagnostic markers for ESCA liquid biopsy. By mutation analysis of cfDNA NGS data, this study finally identified 37 genetically altered ESCA-specific genes. The functional enrichment analysis showed that these genes have close relationships with the occurrence and development of cancer (Fig. S8).

The study was performed with a relatively small sample size at a single institution. This study analyzed 20 cfDNA samples including 4 cfDNA samples from normal people, 4 cfDNA samples from post-operation cancer patients and 12 pre-operation cancer patients, in which one normal cfDNA samples could not be used in the subsequent bioinformatics analysis due to too limited sequencing depth. In the verification study, only ten new cfDNA samples from pre-operation cancer patients were used. Therefore, more normal and post-operation cfDNA samples should be included in the future study for the further validation of the current findings. It would be of value if future studies could be designed to address the clinical value of the detection of cfDNA biomarkers in larger sample cohorts. Additionally, only the cfDNA samples from one kind cancer, ESCA, were investigated. Therefore, this study only identified ESCA-associated markers, whether these markers are ESCA specific should be further investigated by analyzing cfDNA samples from various cancers. However, this more complexed investigation can be effectively performed using the same pipeline, SALP-seq plus machine learning.

## Conclusion

In conclusion, we have successfully analyzed many cfDNA samples from ESCA and normal participants by combining SALP-seq and machine learning, identifying both epigenetic and genetic biomarkers of ESCA. These biomarkers can be used to effectively classify cfDNAs from ESCA patients and normal participants. These biomarkers also shed important new insights on the potential regulatory and molecular mechanisms of tumorigenesis of ESCA. This study thus provides a new pipeline for finding new molecular markers for cancers from cfDNA by combining SALP-seq and machine learning. Finally, this study sheds important new insights on the clinical worth of cfDNA.

## Availability of data and materials

The dataset supporting the conclusion of this article is available in the GEO repository: GSE136541. Supplementary Data are available online.

## Acknowledgements

This work was supported by the National Natural Science Foundation of China (61971122).

